# NeuRiPP: Neural network identification of RiPP precursor peptides

**DOI:** 10.1101/616060

**Authors:** Emmanuel L.C. de los Santos

## Abstract

Significant progress has been made in the past few years on the computational identification biosynthetic gene clusters (BGCs) that encode ribosomally synthesized and post-translationally modified peptides (RiPPs). This is done by identifying both RiPP tailoring enzymes (RTEs) and RiPP precursor peptides (PPs). However, identification of PPs, particularly for novel RiPP classes remains challenging. To address this, machine learning has been used to accurately identify PP sequences. However, current machine learning tools have limitations, since they are specific to the RiPP-class they are trained for, and are context-dependent, requiring information about the surrounding genetic environment of the putative PP sequences. NeuRiPP overcomes these limitations. It does this by leveraging the rich data set of high-confidence putative PP sequences from existing programs, along with experimentally verified PPs from RiPP databases. NeuRiPP uses neural network models that are suitable for peptide classification with weights trained on PP datasets. It is able to identify known PP sequences, and sequences that are likely PPs. When tested on existing RiPP BGC datasets, NeuRiPP is able to identify PP sequences in significantly more putative RiPP clusters than current tools, while maintaining the same HMM hit accuracy. Finally, NeuRiPP was able to successfully identify PP sequences from novel RiPP classes that are recently characterized experimentally, highlighting its utility in complementing existing bioinformatics tools.

## INTRODUCTION

Bacterial specialized metabolites have been a source of bioactive chemical compounds with myriad applications especially in the pharmaceutical and agrochemical industries (1). Advances in DNA sequencing technology and the development of computational tools to identify putative biosynthetic gene clusters (BGCs), have led to a renewed interest in exploring specialized metabolites from microbes as a potential source of novel compounds as sequencing information has suggested that a large fraction of the biosynthetic potential of these microorganisms remains untapped and undetectable under normal laboratory conditions (2, 3). Ribosomally synthesized and post-translationally modified peptides (RiPPs) constitute a diverse class of natural products with a variety of different bioactivities. In contrast to peptide natural products from assembly-line non-ribosomal peptide synthetase (NRPS) pathways, RiPPs are derived from a ribosomally encoded precursor peptide (PP) that is extensively modified by RiPP tailoring enzymes (RTEs) (4, 5). Beginning from a ribosomally encoded peptide makes RiPPs an attractive target for bioengineering as RTEs can be highly selective for recognition sequences in the PP but promiscuously process other regions of the sequence (6). Putative BGCs encoding RiPPs are identified computationally by looking for regions in a genome where there are co-occurences of RTEs and PPs. This makes it relatively easy to identify RiPP BGCs of known RiPP classes by looking for co-localization of RTEs specific to the particular RiPP class. Identification of putative PP sequences is more challenging as they are frequently missed in genome annotation due to their short size (6). However, their proper identification is an important aspect of *in silico* RiPP BGC analysis as knowledge of the PP sequence can aid in structure elucidation and provide information of the molecular interactions between the RTEs and the PP (5). To this end, several methods have been developed to identify putative PPs in regions in proximity to RTEs, this typically involves a two-step process where sequences to be screened are first identified either through the use of gene-finding software (5, 7), or from identifying open reading frames (ORFs) of specified length in the proximity of RTEs (6, 8). The likelihood of these sequences to be PPs is then evaluated by different methods such as looking at similarity to known PPs by BLAST (7), hidden Markov models (HMMs) (5, 9), or machine learning approaches such as Support Vector Machine (SVM) classifiers that are trained to identify likely PPs for different classes based on characteristics of PPs in the specified class (6, 10). While successful in identifying PP sequences and even identifying sequences that are different from known PPs of a specified RiPP class, these approaches have limitations which include only recognizing similar enough sequences to known PPs and being class-specific or context-dependent on the genes surrounding the putative PP. These hinder the development of bioinformatic workflows to identify novel RiPP classes.

One approach to potentially discover novel RiPP classes is to begin by identifying putative PP sequences before exploring the genetic context surrounding the PP sequences for similar sets of RTEs. Due to the large amount of genomic information to process this method requires a context and class-independent way of identifying likely PP sequences. Because there is no genetic window to focus a search, using ORFs to specify the sequences to be classified would result in a large number of sequences and false positives as ORFs do not necessarily correspond to coding sequences particularly in organisms whose GC content is skewed. A recent study presented a pipeline for identifying new RiPP clusters that included a modified version of the gene finding software prodigal (11), prodigal-short. Prodigal-short was used to find putative PPs in proximity to RTEs, and peptide similarity network analysis of the identified PPs was used to identify new RiPP classes (5). This demonstrated the potential of using gene-finding software as a starting point for identifying novel RiPPs; however, the number of likely coding sequences from this approach was still large and the researchers used proximity to known RTEs, restricting searches by phylogeny, and looking at only large similarity networks to reduce the number of putative BGCs to a size where manual curation was tractable. A few of these steps could be avoided if a further context and class-independent step were present to discriminate between likely PPs and false positives. The success of SVM classifiers has led to an increase in the number of high-confidence sequences that are likely to be PPs for several different classes of RiPPs. This along with the increasing number of experimentally verified PP sequences, led me to hypothesize that a positive dataset of reasonable size and quality could be constructed to train a deep neural network (DNN) to classify peptide sequences on their likelihood of being PPs.

DNNs have been successfully employed in image classification problems (12) and text sentiment analysis (13, 14). Problems that could be analogous to peptide classification problems. Neural networks are also gaining popularity in their application to biological systems. Some examples of these in the context of biological sequences are DNNs trained to identify lab origin given a DNA sequence (15), identify whether a sequence of DNA is plasmid or chromosomal in origin (16), and predicting protein-protein interactions between two proteins (17).

In this study, I explore whether DNN architectures successful in image and text classifiers are suitable for the problem of identifying putative PPs. I demonstrate that NeuRiPP, a DNN classifier trained on high confidence PP sequences, is able to provide discriminatory power and enrich for likely PP sequences. NeuRiPP is implemented in Python and is thus easily integratable into existing bioinformatics workflows. It allows for the identification of putative RiPP BGCs starting with the PP instead of the RTEs. The success of the DNN models in discriminating PPs also suggests the suitability of these models for discriminating other types of peptides using the training module of NeuRiPP allowing it to be flexible for other types of peptide classification tasks.

## MATERIALS AND METHODS

### Datasets

#### Positive Set

Positive PP sequences were obtained by collating information from different sources including high-scoring lassopeptide (6) and thiopeptide (18) sequences from RODEO, thiopeptide sequences from Thiofinder (19), lassopeptide, microviridin, and thiopeptide sequences from RiPPER (5), and precursor peptide sequences of various different RiPP classes from PRISM (20). To further supplement the positive set, high scoring lantipeptide, sactipeptide, thiopeptide, and lassopeptide sequences from the antiSMASH database version two were added (21). After dereplication, the final positive set consisted of 2726 sequences. A search consisting of HMMs built to identify known precursor domains as part of Refseq (9) consolidated for RiPPER (5) in addition to further PP HMMs from antiSMASH (22), consisting of a total of 59 different HMMs, was run on the positive set, this resulted in positive HMM hits on 67% of the positive set. A summary of the different HMM PP models can be found in Table S1 in the Supplementary Information.

#### Negative Set

The negative set consisted of low-scoring lassopeptide sequences from the RODEO SVM-classifier (6) and a set of short peptide sequences that were not PPs from Marnix Medema (personal communication), filtered to include only sequences between 20-120 amino acids. This set was collated and checked against the positive set for overlaps. The final negative set consisted of 19224 sequences of which 0.02% were HMM hits.

### Preparation of the Sequence Data as Neural Network Input

A maximum length of 120 amino acids was used as the input for the neural network. Any sequences longer than 120 tested were truncated. Amino acids were represented as a single hot-vector of size 20 where the values in the vector are all 0 except for the amino-acid represented which would have a value of 1 (Figure 1a). Sequences that were less than 120 in length were padded with vectors containing all zeros. This resulted in a uniform input of a 120×20 matrix as the input for the neural network. Positive sequences were tagged with a 1 and negative sequences with a 0. The neural networks were constructed to have a 2×1 output representing the probability that its input is in class 0 or 1 respectively.

**Figure 1.**
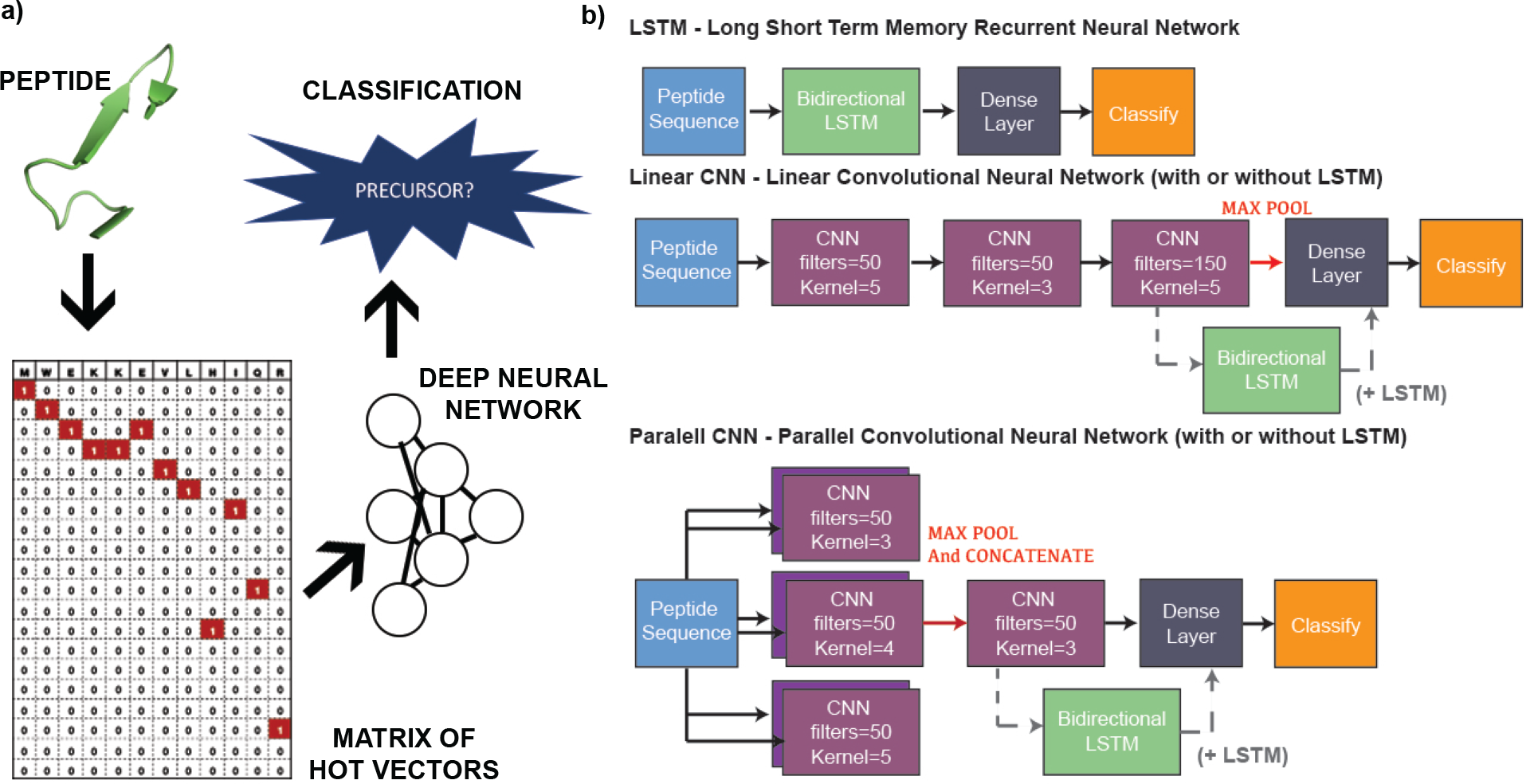
NeuRiPP Workflow and Model Architectures. **(a)** Peptide sequences between 20–120 amino acids long are converted into a 20-by-120 matrix, this serves as an input to a deep neural network with different architectures (see (b)), which determines whether or not the sequence is a likely precursor peptide. **(b)** Five different model architectures were tested, a Long Short-Term Memory Recurrent Neural Network (LSTM), two different Convolutional Neural Network (CNN) layouts, and a combination of the CNN layouts with an LSTM layer.

### Models

Five different DNN architectures were tested. These were inspired by model architectures that were successful in text classification problems (13, 23, 24). All models were implemented in Python 3 using Tensorflow 2.0 (25). To prevent overfitting, a Dropout layer with a 0.5 dropout rate was added between the final densely connected layer and the classification layer. Dropout randomly sets the weights of a set number of nodes (half in this case) to zero preventing the network from becoming reliant on any one node during the training step (26, 27). The last layer of all of the network designs, the classification layer, was a dense layer with a sigmoidal activation function that output a 2 by 1 vector that sums to 1. This could be interpreted as the probability that a given input sequence was a PP. The designs consisted of either long short-term memory recurrent neural network (LSTM) layers, convolutional neural network (CNN) layers, or a combination of both (Figure 1b). Specifically, the five architectures tested were:

- LSTM – Single Bi-directional LSTM with 0.15 dropout and 60 cells. This is followed by a densely connected layer of 60 units and finally, the classification layer.
- Linear CNN – Three successive CNN layers with varying filter and kernel sizes, following the last CNN layer, values are max pooled in groups of 2 before a 40 unit dense layer and the classification layer.
- Parallel CNN – Input is fed in parallel to two CNN layers each of 3 different kernel sizes (6 CNNs total)with max pooling occurring between each of the two layers before concatenating the results. This is fed into a final CNN layer with 150 filters and a kernel size of 3. The output is max pooled in groups of 3 before being fed into a 60 unit Dense Layer and the classification layer.
- Linear CNN + LSTM – Identical to the Linear CNN, with a 60 cell LSTM layer before the dense and classification layer.
- Parallel CNN + LSTM – Identical to the Parallel CNN, with a 60 cell LSTM layer before the dense and classification layer.

### Training

Figure 2 summarizes the procedure used to train the neural network. Because the negative dataset was about 7 times larger than the positive dataset, the negative set was subsampled by 35% in order to prevent over training of the model on negative data. This resulted in a dataset consisting of 9454 sequences per training cycle. 85% of this set was used to train the neural network using sparse cross-entropy as the loss function and adam (28) as the weight optimization algorithm. After weight optimization, the remaining 15% of the dataset was used to test the neural network using total accuracy as the metric. If the round of optimization improved the accuracy of the network, the weights were saved at the end of the training cycle. The negative dataset was resampled every 5 rounds to ensure that the neural network was exposed to the entire negative test set. Weight optimization was halted either if there was no improvement in model accuracy after 50 rounds, or after 200 rounds of training. Final model accuracy was measured on the entire dataset.

**Figure 2.**
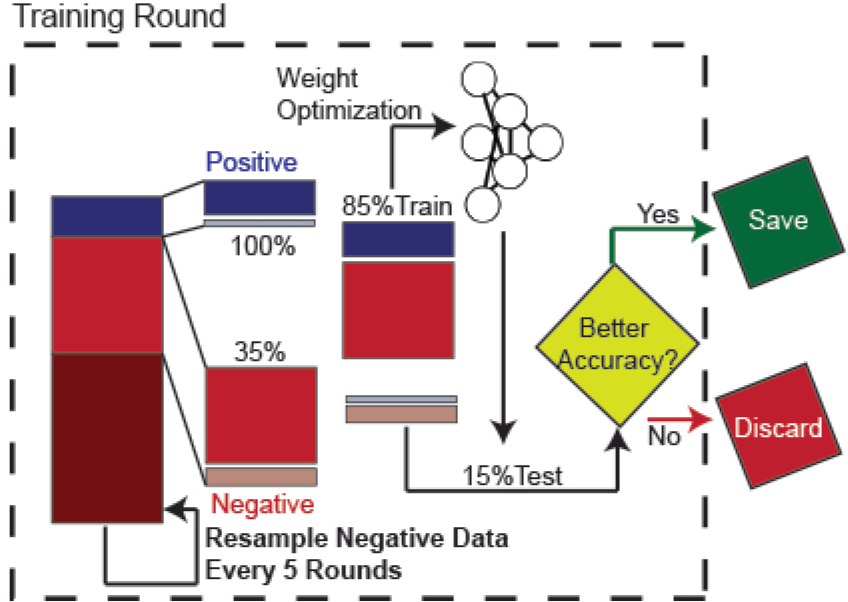
NeuRiPP Training Procedure. Every model architecture was subjected to two hundred rounds of weight optimization. For each round of weight optimization, the entire positive training set, and a randomly subsampled portion of the negative training set is used. 85% of this set is used to optimize the weights of the neural network using adam as an optimizer and cross entropy as the loss function. The remaining 15% was used to test the accuracy of the model. If the weights increased the accuracy of the model, these were stored. The negative set was resampled every five rounds of training.

### Testing

#### antiSMASH database version 2.0

antiSMASH 5 (29) was run on genbank files downloaded from antiSMASH database version 2 (21) corresponding to RiPP and bacteriocin clusters. RODEO precursor peptide predictions were extracted from the json file output of the antiSMASH runs. To obtain candidate sequences for NeuRiPP to classify, prodigal-short (5), a modified version of the prodigal (11, 30) gene finding software was run in “–meta” mode on the fasta sequences of the RiPP clusters identified by antiSMASH from the antiSMASH database. These sequences were classified by the “classify.py” module of NeuRiPP. Comparison to RODEO predictions was done by a custom python script. BGCs were separated into the different RiPP classes using the antiSMASH classification rules (22), or in the case of the thiopeptide class, stricter classification rules using HMMs from a bioinformatic analysis of the thiopeptides (18). If the cluster was identified to be of more than one class by the classification rules, it was counted in both of the RiPP classes.

#### RiPPER Thioviramide predictions

Peptide sequences corresponding to potential thioviramide precursor peptides were obtained from the supplemental information from the “all peptides” section of the RiPPER publication (5). The NeuRiPP classifier was run on the sequences and the predictions were compared to the 30 sequence similarity networks analyzed using a custom python script.

## RESULTS AND DISCUSSION

### NeuRiPP is able to classify peptides in the training set with high accuracy

Table 1 summarizes the best accuracy obtained with for each model architecture on the entire training set. All of the models were able to achieve a high degree of accuracy on the training data. The parallel CNN architecture was the most accurate at 99.84%. In order to check that the high accuracy was not simply to the neural network being overfit to the data (i.e. that the model would only be able to classify peptide sequences it was trained on), the models were also trained on a dataset that randomly excluded 15% of the positive dataset (550 sequences), and 8.6% of the negative set (1650 sequences). The different architectures were trained on the remaining 19750 sequences as previously described. Tables S2 and S3 summarize the accuracy of the architectures on the set of excluded peptides, and the entire training set. When trained with the smaller set, the neural network is less accurate. On the set of sequences that was excluded for training, the LSTM architecture was the most accurate at 98.37% total accuracy. However, when the accuracy was evaluated on the entire training set, the parallel CNN still achieved the highest accuracy at 99.37%. These results suggest that the network is able to capture features in the sequences that are distinct to PPs. The improvements in accuracy when more data are given exhibit the suitability of the tested network architectures for sequence based peptide classification. This is important as NeuRiPP’s performance can be improved as the number of high-confidence PPs increases, as a result of improvements in class-specific PP identifiers, and experimental verification of new RiPPs.

**Table 1.** Accuracy of different network architectures on training set.

| Network | Positive Set Accuracy | Negative Set Accuracy | Total Accuracy |
| --- | --- | --- | --- |
| LSTM | 92.00% | 99.66% | 98.71% |
| Linear CNN | 99.60% | 99.43% | 99.45% |
| Parallel CNN | **99.96%** | **99.82%** | **99.84%** |
| Linear CNN + LSTM | 97.36% | 99.76% | 99.46% |
| Parallel CNN + LSTM | 97.80% | 99.48% | 99.27% |

Given the high accuracy and quick training time (Table S4) of the parallel CNN, this was selected as the default model architecture. The weights that were used were the weights obtained when trained with the entire training set. This was so the network would take advantage of all of the information available when asked to identify putative PPs.

### Sequences identified by NeuRiPP are enriched with HMM hits for known precursor peptides

In order to evaluate NeuRiPP’s role in an existing genome mining workflow, the latest version of antiSMASH (29) was run on sequences from the database. This is a common first step in a genome mining pipeline and ensured that the predictions for RiPP and bacteriocin classes were up to date. This process resulted in 35477 RiPP clusters covering 16 classes of RiPPs (Figure 3a, Table S5)

**Figure 3.**
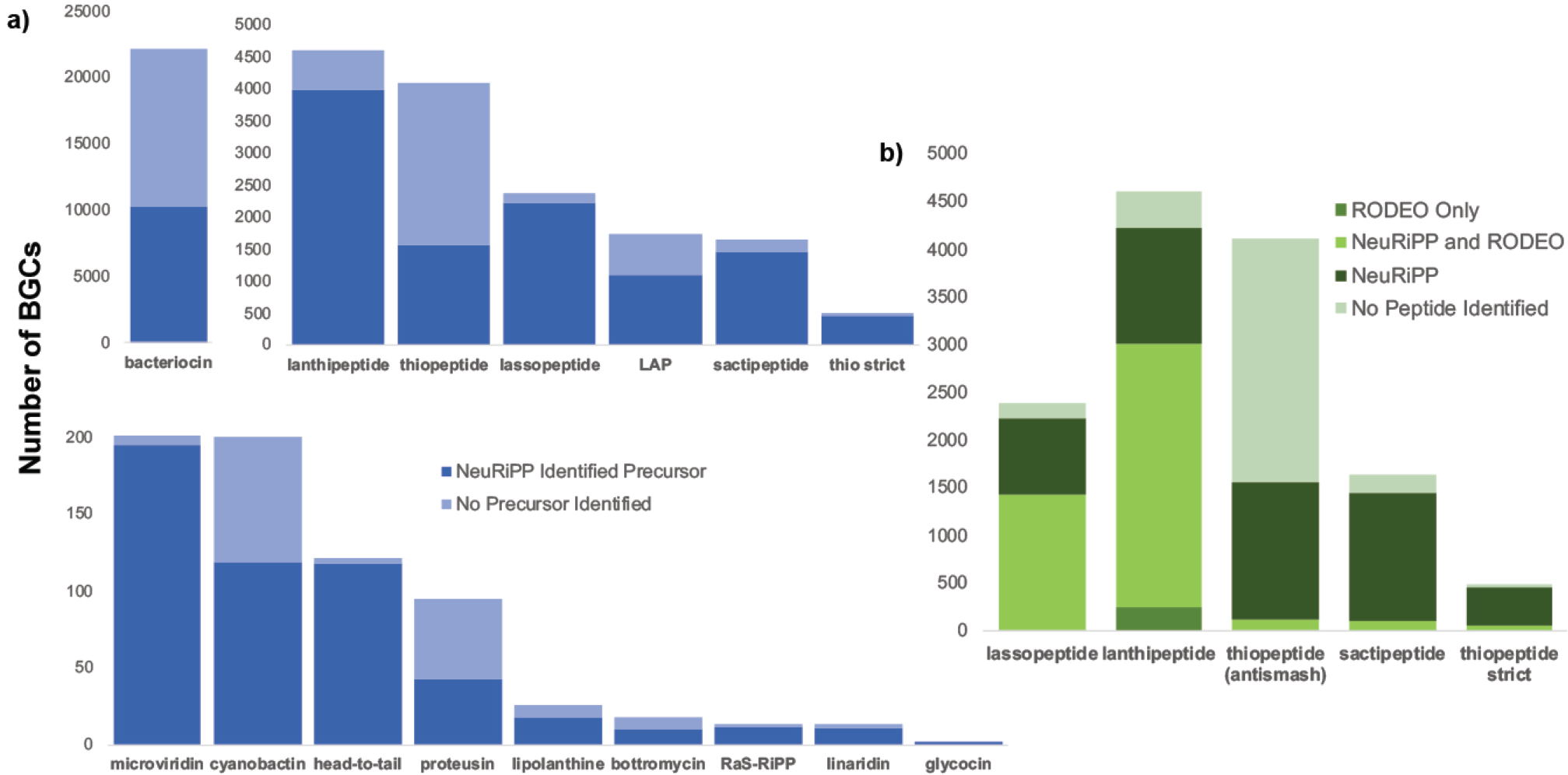
NeuRiPP predictions on RiPP BGCs in the antiSMASH v2 database. **(a)** Breakdown of RiPP clusters in the antiSMASH v2 database by RiPP class. NeuRiPP is able to identify putative PPs in all of the RiPP classes in the antiSMASH database. **(b)** Comparison of NeuRiPP and RODEO Predictions for RODEO-type BGCs. NeuRiPP and RODEO predictions are largely congruent for BGCs where RODEO makes a putative PP prediction. NeuRiPP is able to predict PP sequences in a greater number of RODEO-type RiPP BGCs with a high precursor HMM hit rate on these predictions than RODEO.

Running prodigal-short on these clusters yielded a total of 150366 peptide sequences between 20-120 amino acids long for NeuRiPP to classify. When these sequences were tested on the set of precursor peptide HMMs, 9958 or 6.6% were identified as HMM hits. NeuRiPP classified 34579, or around 20%, of these sequences as putative PPs, with 8485 or 25% of them as HMM hits, a four-fold enrichment from the prodigal-short set. In contrast, there are 1457 (1%) HMM hits on the sequences classified as negatives by NeuRiPP. RODEO identified 8780 peptides as PPs, of which 32% were HMMs hits (Table 2). Figure 3a summarizes the composition of RiPP classes in the database, and whether or not NeuRiPP identified candidates as PPs in the cluster. NeuRiPP makes predictions on PPs in any RiPP BGC regardless of class. It identifies putative PP sequences in 19939 (56.20%) of the RiPP BGCs in the antiSMASH database. Unsurprisingly, it is able to identify putative PPs in a large percentage of clusters in the microviridin, lantipeptide, lassopeptide, and sactipeptide classes as these constitute a large fraction of the classes in the positive training dataset. In the case of thiopeptides, NeuRiPP fails to identify putative PPs in a majority of the thiopeptide clusters in the database. This could be related to the lower accuracy a generic RiPPER search also has in identifying PP in the thiopeptide class (5). It is possible that PPs are more diverse than other RiPP classes, or BGCs classified as thiopeptides are incorrectly classified and actually belong to different RiPP classes that still have not been well-characterized. A bioinformatic study on the thiopeptides that used RODEO to expand the thiopeptide class and discover new thiopeptides developed custom HMMs for the identification of thiopeptide BGCs, using these HMMs to identify thiopeptide BGCs instead of the default antiSMASH detection rules resulted in a smaller subset (489 of 4104 thiopeptide labeled BGCs) being identified as thiopeptide clusters (”thiopeptide strict”). NeuRiPP is able to identify potential PPs in a larger fraction of the strict thiopeptide set. Encouragingly, NeuRiPP is also able to identify potential PP sequences in other RiPP classes. This presents an opportunity of improvement for NeuRiPP as training with a richer, more diverse set of positive PP sequences from different classes could improve overall performance in general and allow it to identify even more putative PP in uncharacterized RiPP classes.

**Table 2.** Summary of Precursor Peptide HMM Hits from Different Classifiers

| Set | Sequences | HMM Hits | % of Set |
| --- | --- | --- | --- |
| Prodigal-short | 150366 | 9958 | 6.62% |
| RODEO Predictions | 8780 | 2773 | 31.58% |
| NeuRiPP Predictions | 34579 | 8485 | 24.54% |
| NeuRiPP (RODEO-type clusters) | 15403 | 5553 | 36.05% |
| NeuRiPP Negative | 115537 | 1474 | 1.28% |

### NeuRiPP predictions complement RODEO predictions for RODEO-type clusters in the antiSMASH database

The antiSMASH database contains 12741 clusters that are classified at as lanthipeptides, sactipeptides, thiopeptides, or lassopeptides. These are the RiPP cluster types that RODEO contains SVMs to identify PPs in. RODEO identifies 8780 peptides (6058 unique sequences) in 4681 RODEO-type clusters as putative PPs. NeuRiPP identifies putative PPs in 4415 (94.32%) of these clusters. When looking at the actual peptide predictions, 5180 (3134 unique, 52%) of the RODEO predicted PPs are identified by NeuRiPP as putative PPs (Table S6). This was partly because a portion of the ORFs (2610 sequences, 30%) that were classified by RODEO were not identified by prodigal-short to be coding sequences, a potential limitation of using gene finding software. However, NeuRiPP is still able to identify potential PP sequences in a majority of the clusters where RODEO identifies PPs. In 4084 (92.05%) of these clusters there is at least one peptide that matches between the NeuRiPP and RODEO prediction. In 3330 (71.14%) of the clusters, all of the RODEO predictions correspond to NeuRiPP hits and in 2142 (45.76%) of the clusters, the RODEO and NeuRiPP predictions match exactly. In 331 clusters, NeuRiPP predicts a different set of sequences as the PPs for the cluster.

It is important to note that a portion (35%) of the RODEO PP sequences that had high RODEO scores in the antiSMASH database were thus used as part of the positive training set; however, NeuRiPP identified an additional 2205 PPs that it was not trained on. NeuRiPP is also able to identify putative PPs in 12475 (98%) of the RODEO-type clusters in the antiSMASH database, while maintaining a relatively high HMM-hit rate of 36% (Table 2), compared to the RODEO predictions. Taken together, these show that NeuRiPP is able to provide additional discriminatory power in identifying putative PPs. At worst, NeuRiPP is able to complement RODEO predictions in a computational pipeline, sequences predicted by both NeuRiPP and RODEO can be accepted with a higher confidence. However, in clusters where RODEO is unable to make prediction, the relatively high HMM hit rate suggests that the PP predictions that NeuRiPP makes on its own can be taken as potential PP sequences. Figure 3b summarizes the NeuRiPP and RODEO predictions in the RODEO-type clusters in the antiSMASH database. Interestingly, neither NeuRiPP nor RODEO are able to provide PP predictions for a majority of the thiopeptide clusters when classified using the default antiSMASH rules, this discrepancy is resolved when stricter classification rules are used (”thiopeptide strict”) (18), highlighting the need for further characterization and classification even in known RiPP classes.

### NeuRiPP identifies novel thioamidated peptides identified by RiPPER

In order to identify new families of thioamidated peptides, researchers who developed the RiPPER methodology employed it to analyze regions of DNA in proximity to co-occurences of a YcaO-domain containing protein and a TfuA-like protein in *Streptomyces* genomes (5). RiPPER retrieved 743 peptides which were further analyzed using peptide similarity networking. The genetic environment surrounding the thirty peptide similarity networks containing at least four sequences was examined for gene conservation and Pfam (31) domain composition in order to determine whether or not the similarity networks represented likely precursor peptides. With this analysis, they labeled twelve of the peptide similarity networks PP networks as “yes” concerning whether or not they contained likely PPs. These included the peptide similarity networks that contained thioviramide, a known thioamidated RiPP, and thiovirsolin, a novel thioamidated RiPP that was part of a new thioamidated RiPP family, predicted using the RiPPER workflow, purified and characterized. Five of the remaining peptide similarity networks were labeled “maybe” as likely precursor peptides. To further demonstrate NeuRiPP’s utility in a genome mining pipeline for discovering novel RiPPs, the 743 peptides retrieved using RiPPER for creating the thioamidated PP similarity networks were classified by NeuRiPP. Unlike the training set, NeuRiPP had been previously unexposed to these sequences, with the exception of the thioviramide sequence that was included in PRISM (20) and a second sequence that was previously identified by Thiofinder as a thiopeptide precursor (19). NeuRiPP identified 91 of these sequences as likely PPs. Eight of the thioamidated peptide similarity networks analyzed in RiPPER contained multiple NeuRiPP hits (Table S7), these included the similarity networks that contained thioviramide (Network 5) and thiovarsolin (Network 22). Seven of these networks were determined to be likely precursor peptide sequences, while the other network that contained multiple NeuRiPP hits was thought to be possibly a PP sequence network. While NeuRiPP did not have multiple hits in the other five similarity networks that were thought to be likely precursor peptide sequences, there is a much greater chance that a similarity network containing multiple NeuRiPP hits is a likely PP. This suggests that a workflow where sequences extracted by RiPPER can first be classified by NeuRiPP before generating the peptide similarity networks will be enriched for likely PPs and RiPP BGCs. This is beneficial as it reduces the amount of clusters that have to be manually examined and further analyzed. Only 12% of the sequences obtained by RiPPER were NeuRiPP hits, while maintaining a high rate of discovering RiPP BGCs from the peptide similarity networking analysis.

## CONCLUSION

NeuRiPP is a fast, easy to use tool that is able to predict putative PP sequences in a class-independent manner. It is able to complement existing RiPP bioinformatics tools by either confirming their predictions, or offering predictions in BGCs where other tools are unable to make predictions. Peptide sequences classified as NeuRiPP hits show a similar or higher HMM hit rate to precursor peptide HMMs in existing tools. NeuRiPP is easily integratable into the antiSMASH and RODEO workflows. It also fits well with the RiPPER methodology by adding an additional filtering step before peptide similarity networking, reducing the amount of manual analysis and curation that needs to be done, while maintaining a high hit of likely PP sequences.

The increased selectivity and discrimination of NeuRiPP along with its class-independence also allow for a RiPP mining methodology that is independent of preliminary knowledge of RTEs allowing for the discovery of novel RiPP classes. Most existing bioinformatics tools for RiPP mining, such as BAGEL (8), RODEO (32), antiSMASH (29), RiPP Miner (10) and PRISM (20), provide a wealth of information on well-characterized RiPP classes by identifying putative RiPP BGCs based on the set of conserved protein domains responsible for the biosynthesis of the specific RiPP class. They provide further information often predicting the mass, cleavage sites, potential modifications, and the sequence of mature peptide. RiPPER (5) works well as a complement to these tools by allowing the user to specify the gene clusters and types to be examined. However, there is still a large number of sequences retrieved by RiPPER based on prodigal-short, requiring some sort of filtering step that reduces the number of sequences to examine. This is often dependent on prior knowledge about a specific class of enzymes that could potentially be involved in RiPP-biosynthesis which biases searches towards “known unknowns”. By providing an additional filtering step, NeuRiPP can potentially overcome this allowing peptide and gene similarity networks to be constructed without first having to specify a search space with a specific enzyme or domain as a seed. While NeuRiPP is limited in the fact that it will be biased towards the precursor peptide classes in its training set, having multiple RiPP classes as exemplars can potentially overcome some of these biases allowing the neural network to discern common characteristics in PP sequences across different RiPP classes. By not starting with RTEs as seeds for the search, NeuRiPP can potentially identify BGCs that contain novel combinations of known RTEs from the classes it was trained on, or potentially even completely new sets of RTEs.

Finally, the neural network structure of NeuRiPP allows for flexibility and offers to potential for further improvements. NeuRiPP model weights can be retrained to improve its performance on a specific RiPP class that is of particular interest. As more RiPP classes are discovered and experimentally verified, PP sequences from these can be added to the positive training set which should improve NeuRiPP’s general performance. Training weights for the NeuRiPP models are fast and do not require intensive computing power (Table S4). The optimized weights of the the parallel CNN model used in this study were trained on a laptop computer in a few hours. While NeuRiPP was trained on PPs, the neural network architecture may be suitable for other peptide classification problems. The training module is flexible and only requires fasta files of positive and negative examples of amino acid sequences, allowing the possible extension of NeuRiPP as a general protein classifier.

## AVAILABILITY

NeuRiPP is available at: https://github.com/emzodls/neuripp under the GNU AGPL v3. The repository contains the training sets described in the study, along with the optimized weights for each of the model architectures. The train module can be used to create a custom set of model weights, while the classify module can be used with the pre-trained weights or new weights to identify PP sequences.

## ACKNOWLEDGEMENTS

ELCdlS is a Research Career Development Fellow in the Warwick Integrative Synthetic Biology Centre, which is supported by a grant from the BBSRC and EPSRC (BB/M017982/1). Marnix Medema for providing negative sequences used to train NeuRiPP. Andrew Truman, Anandh Swaminatham, Bryce Kelly, Alexander Kloosterman, Marnix Medema, and Douglas Mitchell for discussions about NeuRiPP.

